# The effect of carbon subsidies on marine planktonic niche partitioning and recruitment during biofilm assembly

**DOI:** 10.1101/013938

**Authors:** Charles Pepe-Ranney, Edward Hall

**Affiliations:** Cornell University, Department of Crop and Soil Sciences, Ithaca, NY, USA; Colorado State University, Natural Resource and Ecology Laboratory, Fort Collings, CO, USA

**Keywords:** microbial ecology, 16S, 23S, planktonic, biofilm, carbon subsidies, resource stoichiometry

## Abstract

The influence of resource availability on planktonic and biofilm microbial community membership is poorly understood. Heterotrophic bacteria derive some to all of their organic carbon (C) from photoautotrophs while simultaneously competing with photoautotrophs for inorganic nutrients such as phosphorus (P) or nitrogen (N). Therefore, C inputs have the potential to shift the competitive balance of aquatic microbial communities by increasing the resource space available to heterotrophs (more C) while decreasing the resource space available to photoautotrophs (less mineral nutrients due to increased competition from heterotrophs). To test how resource dynamics affect membership of planktonic communities and assembly of biofilm communities we amended a series of flow-through mesocosms with C to alter the availability of C among treatments. Each mesocosm was fed with unfiltered seawater and incubated with sterilized microscope slides as surfaces for biofilm formation. The highest C treatment had the highest planktonic heterotroph abundance, lowest planktonic photoautotroph abundance, and highest biofilm biomass. We surveyed bacterial 16S rRNA genes and plastid 23S rRNA genes to characterize biofilm and planktonic community membership and structure. Regardless of resource additions, biofilm communities had higher alpha diversity than planktonic communities in all mesocosms. Heterotrophic plankton communities were distinct from heterotrophic biofilm communities in all but the highest C treatment where heterotrophic plankton and biofilm communities resembled each other after 17 days. Unlike the heterotrophs, photoautotrophic plankton communities were different than photoautotrophic biofilm communities in composition in all treatments including the highest C treatment. Our results suggest that although resource amendments affect community membership and structure, microbial lifestyle (biofilm versus planktonic) has a stronger influence on community composition.

## 1 INTRODUCTION

Biofilms are diverse and complex microbial consortia, and, the biofilm lifestyle is the rule rather than the exception for microbes in many environments. Large and small-scale biofilms architectural features play an important role in their ecology and influence their role in biogeochemical cycles (**Battin et al.**, 2007). Fluid mechanics impact biofilm structure and assembly (**Hodl et al.**, 2011; **Besemer et al.**, 2009; **Battin et al.**, 2003), but it is less clear how other abiotic factors such as resource availability affect biofilm assembly. Aquatic biofilms initiate with seed propagules from the planktonic community (**Hodl et al.**, 2011; **McDougald et al.**, 2011). Thus, resource amendments that influence planktonic communities may also influence the recruitment of microbial populations during biofilm community assembly.

In a crude sense, biofilm and planktonic microbial communities divide into two key groups: oxygenic phototrophs including eukaryotes and cyanobacteria (hereafter “photoautotrophs”), and heterotrophic bacteria and archaea. This dichotomy, admittedly an abstraction (e.g. non-phototrophs can also be autotrophs), can be a powerful paradigm for understanding community shifts across ecosystems of varying trophic state (**Cotner and Biddanda**, 2002). Heterotrophs meet some to all of their organic carbon (C) requirements from photoautotroph produced C while simultaneously competing with photoautotrophs for limiting nutrients such as phosphorous (P) (**Bratbak and Thingstad**, 1985). The presence of external C inputs, such as terrigenous C leaching from the watershed (**Jansson et al.**, 2008; **Karlsson et al.**, 2012) or C exudates derived from macrophytes (**Stets and Cotner**, 2008a,b), can alleviate heterotroph reliance on photoautotroph derived C and shift the heterotroph-photoautotroph relationship from commensal and competitive to strictly competitive (see **Stets and Cotner**, 2008a, Figure 1). Therefore, increased C supply should increase the resource space available to heterotrophs and increase competition for mineral nutrients decreasing nutrients available for photoautotrophs (assuming that heterotrophs are superior competitors for limiting nutrients as has been observed (see **Cotner and Wetzel**, 1992, Figure 1)). These dynamics should result in the increase in heterotroph biomass relative to the photoautotroph biomass along a gradient of increasing labile C inputs. We refer to this differential allocation of limiting resources among components of the microbial community as niche partitioning.

**Figure 1.**
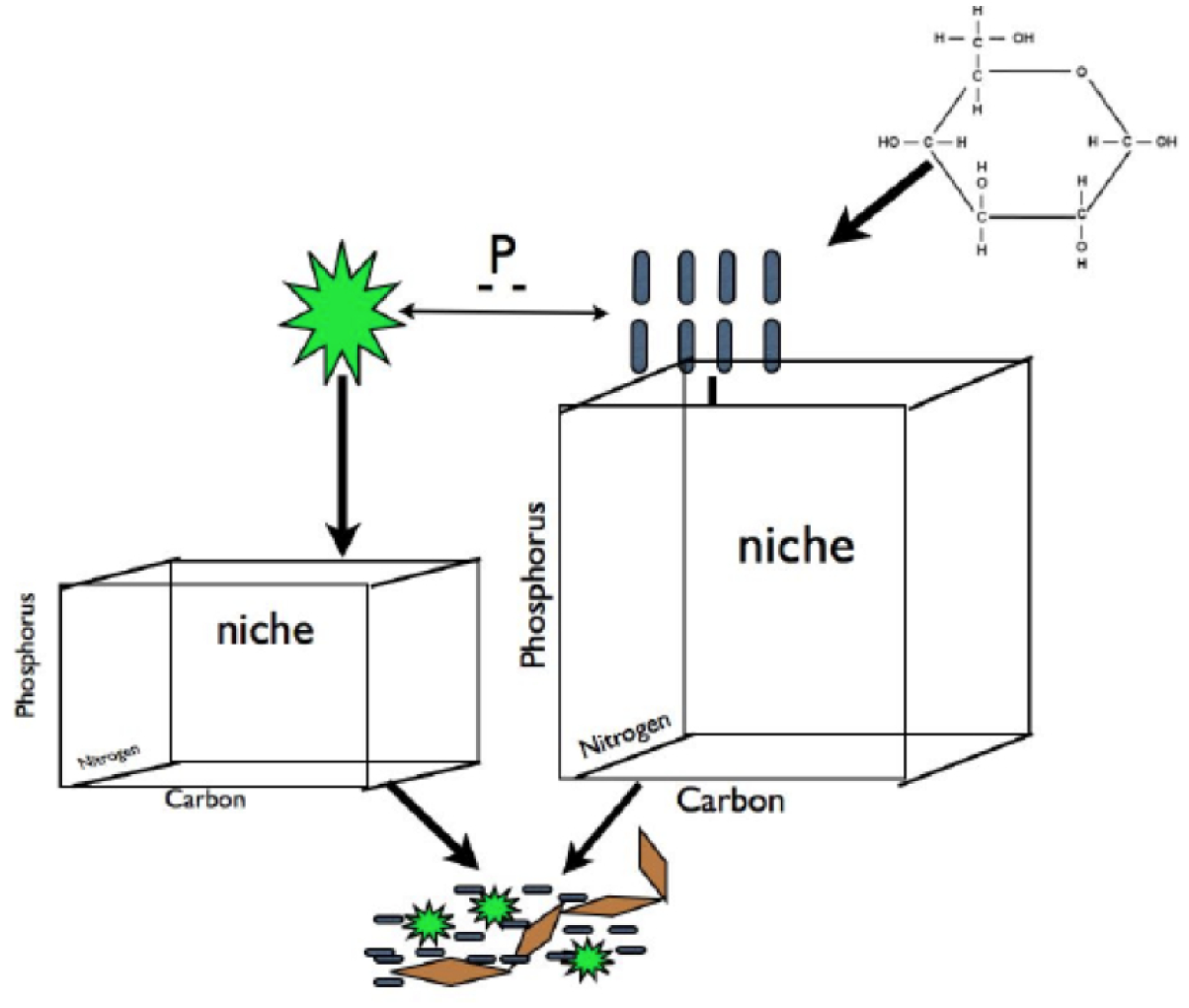
Carbon subsidies in the form of glucose alleviate the dependence of heterotrophic bacteria on photoautotroph derived C exudates. This should result in an increase in resource space and biomass for heterotrophs and a decrease in resource space and biomass for photoautotrophs due to increased competition for mineral nutrients (for simplicity we illustrate competition for P but this is equally applicable other elements that may limit primary production). We hypothesized that this predicted change in biomass pool size of these two groups will result in changes in the plankton community composition of both groups that will propagate to to the composition of biofilm communities for both groups. We refer to shifts in the demand and availability of resources among components of the microbial community as ’partitioning. Blue rods represent heterotrophs, green stars represent photoautotrophs, brown diamonds represent EPS or other cohesive components of a biofilm.

While these gross level dynamics have been discussed conceptually (**Cotner and Biddanda**, 2002) and to some extent demonstrated empirically (**Stets and Cotner**, 2008a), the effects of biomass dynamics on photoautotroph and heterotroph membership and structure has not been directly evaluated in plankton or biofilms. In addition, how changes in planktonic communities propagate to biofilms during community assembly is not well understood. We designed this study to test if C subsidies shift the biomass balance between autotrophs and heterotrophs within the biofilm or its seed pool (i.e. the plankton) and, to measure how changes in biomass pool size alter composition of the plankton and biofilm communities. Specifically, we amended marine mesocosms with varying levels of labile C input and evaluated differences in photoautotroph and heterotrophic bacterial biomass in plankton and biofilm samples along the C gradient. In each treatment we characterized plankton and biofilm community composition by PCR amplifying and DNA sequencing 16S rRNA genes and plastid 23S rRNA genes.

## 2 MATERIALS AND METHODS

### 2.0.1 Experimental Design

Test tube racks were placed in one smaller (185 L, control) and 3 larger (370 L) flow-through mesocosms. All mesocosms were fed directly with marine water from an inflow source in Great Bay, Woods Hole, MA, approximately 200 m from the shore. Each mesocosm had an adjustable flow rate that resulted in a residence time of approximately 12 h. Irregular variation in inflow rate caused the actual mesocosm flow rate to vary around the targeted flow rate throughout the day. However, regular monitoring ensured that the entire volume of each system was flushed approximately two times per day (i.e. maintained a residence time of ∼12 h). To provide a surface for biofilm formation, we attached coverslips to glass slides using nail polish and then attached each slide to the test tube racks using office-style binder clips. Twice daily 10 mL of 37 mM KPO_4_ and 1, 5 and 50 mL of 3.7 M glucose were added to each of 3 mesocosms to achieve target C:P resource amendments of 10, 100 and 500. The goal of the resource amendments was to create a gradient of labile carbon. The same amount of P was added to each treated mesocosm to ensure that response to additions of C were not inhibited by extreme P limitation. The control mesocosm did not receive any C or P amendments.

### 2.0.2 DOC and Chlorophyll Measurements

To assess the efficacy of the C additions, we sampled each mesocosm twice daily during the first week of the experiment to evaluate dissolved organic C (DOC) content. After the initiation of the experiment we collected plankton on filters regularly to evaluate planktonic Chl *a* and bacterial abundance. Once it was clear that pool size of each community had been altered (day 8) we filtered plankton onto 0.2 *μ*m filters and harvested coverslips to assess bacterial and algal biofilm community composition (16S and plastid 23S rDNA). In addition, all mesocosms were analyzed for community composition a second time (day 17) to assess how community composition of both the plankton and biofilm communities had changed over time. Control samples were only analyzed for community composition on day 17.

Samples for dissolved organic C (DOC) analysis were collected in acid washed 50 mL falcon tubes after filtration through a 0.2 *μ*m polycarbonate membrane filter (Millipore GTTP GTTP02500, Sigma Aldrich P9199) attached to a 60 mL syringe. Syringes and filters were first flushed multiple times with the control sample to prevent leaching of C from the syringe or the filter into the sample. Samples were then frozen and analyzed for organic C content with a Shimadzu 500 TOC analyzer (**Wetzel and Likens**, 2000). Biomass of all biofilm samples were measured by difference in pre-(without biofilm) and post-(with biofilm) weighed GF/F filters after oven drying overnight at 60 °C.

For Chl *a* analysis, we collected plankton on GF/F filters (Whatman, Sigma Aldrich Cat. # Z242489) by filtering between 500 mL and 1L from the water column of each mesocosm for each treatment. For biofilm samples, all biofilm was gently removed from the complete area of each coverslip (3 coverslips for each treatment per sampling event) before being placed in a test tube for extraction with 90-95% acetone for ∼32 hours at −20 °C and analyzed immediately using a Turner 10-AU fluorometer (**Wetzel and Likens**, 2000).

Bacterial abundance of the planktonic samples was analyzed using DAPI staining and direct visualization on a Zeis Axio epifluorescence microscope after the methods of Porter and Feig (1980). Briefly, 1-3 mL of water was filtered from three separate water column samples through a 0.2 *μ*m black polycarbonate membrane filter and post stained with a combination of DAPI and Citifluor mountant media (Ted Pella Redding, Ca) to a final concentration of 1 *μ*L mL^−1^.

### 2.0.3 DNA extraction

For plankton, cells were collected by filtering between 20 – 30 mL of water onto a 0.2 *μ*m pore-size polycarbonate filter (Whatman Nucleopore 28417598, Sigma-Aldrich cat# WHA110656). For biofilm communities, biomass from the entire coverslip area of three separate slides was collected and combined in an Eppendorf tube by gently scraping the slip surface with an ethanol rinsed and flamed razor blade. DNA from both the filter and the biofilm was extracted using a Mobio Power Soil DNA isolation kit (MoBio Cat. # 12888).

### 2.0.4 PCR

Samples were amplified for 454 sequencing using a forward and reverse fusion primer. The forward primer was constructed with (5’-3’) the Roche A linker, an 8-10 bp barcode, and the forward gene specific primer sequence. The reverse fusion primer was constructed with (5’-3’) a biotin molecule, the Roche B linker and the reverse gene specific primer sequence. The gene specific primer pair for bacterial SSU rRNA genes was 27F/519R (**Lane**, 1991). The primer pair p23SrV f1/p23SrV r1 was used to target 23S rRNA genes on plastid genomes (**Sherwood and Presting**, 2007). Amplifications were performed in 25 *μ*L reactions with Qiagen HotStar Taq master mix (Qiagen Inc, Valencia, California), 1 *μ*L of each 5 uM primer, and 1 *μ*L of template. Reactions were performed on ABI Veriti thermocyclers (Applied Biosytems, Carlsbad, California) under the following thermal profile: 95°C for 5 min, then 35 cycles of 94°C for 30 sec, 54°C for 40 sec, 72°C for 1 min, followed by one cycle of 72°C for 10 min and 4°C hold. Amplification products were visualized with eGels (Life Technologies, Grand Island, New York). Products were then pooled equimolar and each pool was cleaned with Diffinity RapidTip (Diffinity Genomics, West Henrietta, New York), and size selected using Agencourt AMPure XP (BeckmanCoulter, Indianapolis, Indiana) following Roche 454 protocols (454 Life Sciences, Branford, Connecticut). Size selected pools were then quantified and 150 ng of DNA was hybridized to Dynabeads M-270 (Life Technologies) to create single stranded DNA following Roche 454 protocols (454 Life Sciences). Single stranded DNA was diluted and used in emPCR reactions, which were performed and subsequently enriched. Sequencing followed established manufacture protocols (454 Life Sciences).

### 2.0.5 Sequence quality control

The 16S rRNA gene and plastid 23S rRNA gene sequence collections were demultiplexed and sequences with sample barcodes not matching expected barcodes were discarded. We used the maximum expected error metric (**Edgar**, 2013) calculated from sequence quality scores to cull poor quality sequences from the dataset. Specifically, we discarded any sequence with a maximum expected error count greater than 1 after truncating the sequence to 175 nt. The forward primer and barcode was trimmed from the remaining reads. We checked that all primer trimmed, error screened, and truncated sequences were derived from the targeted region of the LSU or SSU rRNA gene (23S and 16S sequences, respectively) by aligning the reads to Silva LSU or SSU rRNA gene alignment (“Ref” collection, release 115) with the Mothur (**Schloss et al.**, 2009) NAST-algorithm (**DeSantis et al.**, 2006) aligner and inspecting the alignment coordinates. Reads falling outside the expected alignment coordinates were culled from the dataset. Remaining reads were trimmed to consistent alignment coordinates such that all reads began and ended at the same position in the SSU rRNA gene and screened for chimeras with UChime in “denovo” mode (**Edgar et al.**, 2011) via the Mothur UChime wrapper. 19,978 of 56,322 16S rRNA gene sequencing reads and 44,719 or 78,695 plastid 23S rRNA gene sequencing reads passed quality control.

### 2.0.6 Taxonomic annotations

Sequences were taxonomically classified using the UClust (**Edgar**, 2010) based classifier in the QIIME package (**Caporaso et al.**, 2010) with the Greengenes database and taxonomic nomenclature (version “gg_13_5” provided by QIIME developers, 97% OTU representative sequences and corresponding taxonomic annotations, (**McDonald et al.**, 2012)) for 16S reads or the Silva LSU database (“Ref” set, version 115, EMBL taxonomic annotations, (**Quast et al.**, 2013)) for the 23S reads as reference. We used the default parameters for the algorithm (i.e. minimum consensus of 51% at any rank, minimum sequence identity for hits at 90% and the maximum accepted hits value was set to 3).

### 2.0.7 Sequence clustering

Reads were clustered into OTUs following the UParse pipeline. Specifically USearch (version 7.0.1001) was used to establish cluster centroids at a 97% sequence identity level from the quality controlled data and to map quality controlled reads to the centroids. The initial centroid establishment algorithm incorporates a quality control step wherein potentially chimeric reads are not allowed to become cluster seeds. Additionally, we discarded singleton reads. Eighty-eight and 98% of quality controlled reads could be mapped back to our cluster seeds at a 97% identity cutoff for the 16S and 23S sequences, respectively.

### 2.0.8 Alpha and Beta diversity analyses

Alpha diversity calculations were made using PyCogent Python bioinformatics modules (**Knight et al.**, 2007). Rarefaction curves show average OTU counts from 25 re-samplings at intervals of 10 sequences for each sample. Beta diversity analyses were made using Phyloseq (**McMurdie and Holmes**, 2014) and its dependencies (**Oksanen et al.**, 2013). A sparsity threshold of 25% was used for ordination of both plastid 23S and bacterial 16S libraries. Additionally, we discarded any OTUs from the plastid 23S rRNA gene data that could not be annotated as belonging in the Eukaryota or cyanobacteria for differential abundance, ordination and Adonis analyses. Cyanobacterial DNA sequences were removed from 16S rRNA gene sequence collections for ordination, Adonis and differential abundance analyses. We operated under the assumption that non-cyanobacterial bacteria are predominantly heterotrophs in our mesocosm setup and refer to non-cyanobacterial bacteria as “heterotrophs” in the manuscript (this abstraction is useful however there are likely exceptions – i.e. autotrophs among the non-cyanobacterial bacteria). All DNA sequence based results were visualized using GGPlot2 (**Wickham**, 2009). Adonis tests and principal coordinate ordinations were performed using the Bray-Curtis similarity measure for pairwise library comparisons. Adonis tests employed the default value for number of permutations (999) (”adonis” function in Vegan R package, **Oksanen et al.** (2013)). Principal coordinates of OTUs were found by averaging site principal coordinate values for each OTU with OTU relative abundance values (within sites) as weights. The principal coordinate OTU weighted averages were then expanded to match the site-wise variances (**Oksanen et al.**, 2013).

### 2.0.9 Identifying enriched OTUs

We used an RNA-Seq differential expression statistical framework to find OTUs enriched in the given sample classes (R package DESeq2 developed by **Love et al.** (2014)) (for review of RNA-Seq differential expression statistics applied to microbiome OTU count data see **McMurdie and Holmes** (2014)). We use the term differential abundance coined by **McMurdie and Holmes** (2014) to denote OTUs that have different relative abundance across sample classes. We were particularly interested in two sample classes: 1) lifestyle (biofilm or planktonic) and, 2) high C (C:P = 500) versus not high C (C:P = 10, C:P = 100 and C:P = control). A differentially abundant OTU is enriched on one side of a sample class (e.g. enriched in the biofilm versus the plankton). Differential abundance could mark an enrichment of the OTU towards either side of the sample class and the direction of the enrichment is apparent in the sign (positive or negative) of the enrichment. Differential abundance was moderated (see **Love et al.** (2014)) such that the fold change OTU of an OTU across two categories of a sample class can be used to rank the enrichment of OTUs that span a wide range of base abundance. The DESeq2 RNA-Seq statistical framework has been shown to improve power and specificity when identifying differentially abundant OTUs across sample classes in microbiome experiments **McMurdie and Holmes** (2014).

The specific DESeq2 (**Love et al.**, 2014) parameters we used were as follows: All dispersion estimates from DESeq2 were calculated using a local fit for mean-dispersion. Native DESeq2 independent filtering was disabled in favor of explicit independent filtering by sparsity. The sparsity thresholds that produced the maximum number of OTUs with adjusted p-values for differential abundance below a false discovery rate of 10% were selected. Cook’s distance filtering was also disabled when calculating p-values with DESeq2. We used the Benjamini-Hochberg method to adjust p-values for multiple testing (**Benjamini and Hochberg**, 1995). Identical DESeq2 methods were used to assess enriched OTUs from relative abundances grouped into high C (C:P = 500) or low C (C:P *<* 500 and control) categories.

IPython Notebooks with computational methods used to create all figures and tables as well as taking raw sequences through quality control preprocessing are provided at the following URL: https://github.com/chuckpr/BvP_manuscript_figures.

Version information for all R libraries is provided at the end of each IPython Notebook.

## 3 RESULTS

### 3.1 BULK COMMUNITY CHARACTERISTICS

We first assessed the effect of the resource treatments on the dissolved chemistry and bulk community characteristics of the plankton and the biofilms. In the control treatment the mean DOC level was 0.12 +/-0.02 *μ*moles L^−1^. The lower C treatment (C:P = 10) was statistically indistinguishable from the control at 0.10 +/- 0.02 *μ*moles C L^−1^. The intermediate treatment (C:P = 100) increased in DOC to 0.70 *μ*moles C L^−1^ before decreasing to 0.12 *μ*moles C L^−1^ at the end of the experiment, with a mean of 0.28 +/- 0.16 *μ*moles C L^−1^ over the course of the experiment. Only the highest C treatment (C:P=500) had DOC levels that were significantly higher (2.53 +/- 1.6 *μ*moles C L^−1^) than the control treatment, over the course of the experiment. The high DOC levels in the highest C treatment were consistent with C being supplied in excess of the metabolic requirements of the community (i.e. C saturation), but not higher than what has been observed in coastal marine ecosystems.

This increase in DOC in the higher C treatments was associated with decreases in planktonic Chl *a* in each treatment (Figure 2A), however there was no significant difference in biofilm Chl *a* among treatments (Figure 2B). In combination with the decrease in planktonic Chl *a* on the 6th day of the experiment the highest C treatment had approximately 4-fold higher planktonic heterotroph abundance than the control and the 10 *μ*M C treatment (Figure 2D). Similarly, biofilms had significantly higher total biomass in the high C treatment compared to the other treatments (Figure 2D). Thus the shift in resource C:P altered the pool size of both the photoautotroph and heterotroph communities. Clear differences in heterotroph and photoautotroph pool size among treatments allowed us to address how shifts in pool sizes were related to community membership and structure within and among plankton and biofilm communities.

**Figure 2.**
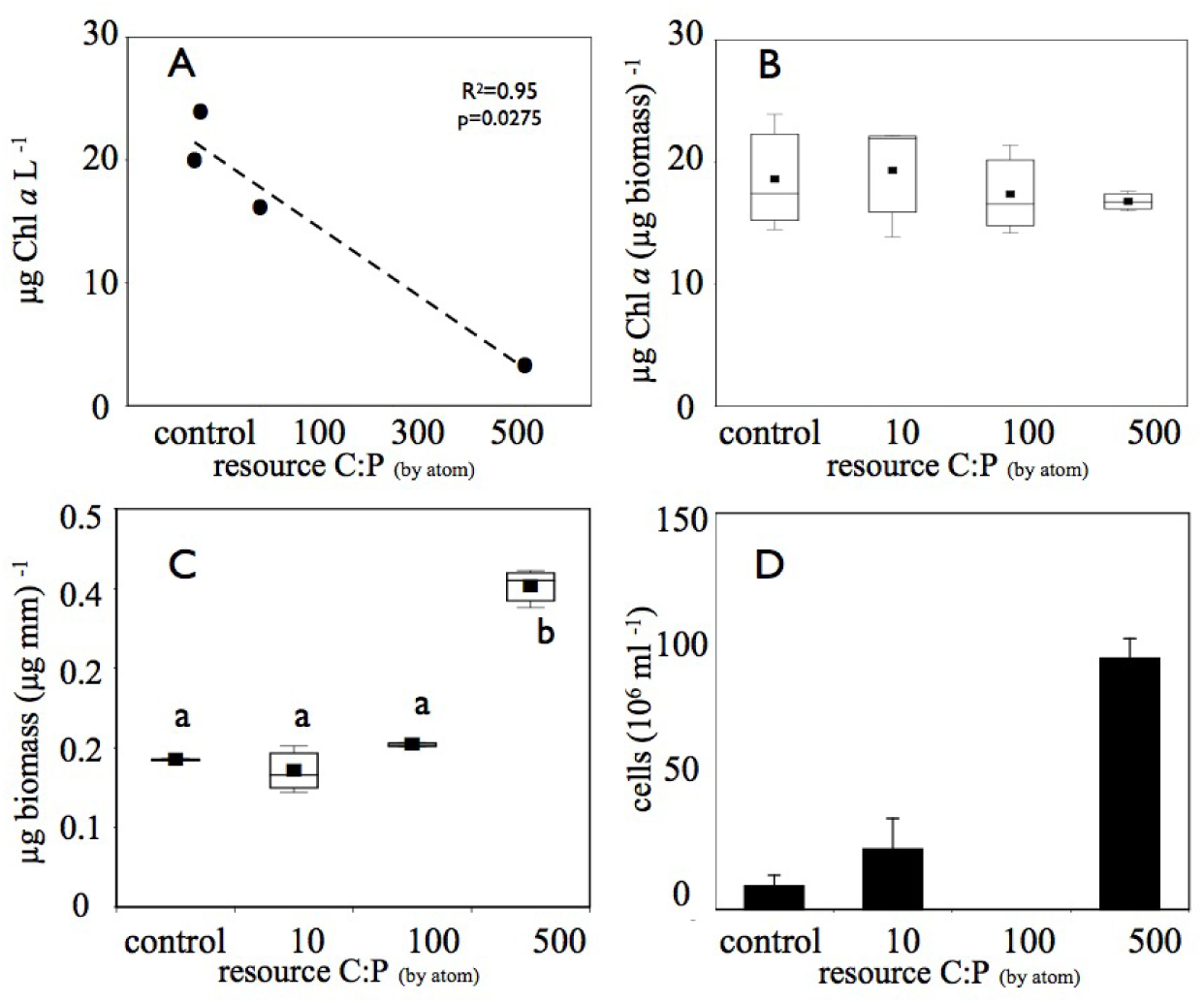
Increased C amendments diminished A) planktonic photoautotroph biomass (estimated as Chl *a*) but B) not biofilm photoautotroph biomass. In contrast, both C) biofilm total biomass and D) number of planktonic bacterial cells increased with increasing C subsidies. Only the highest C treatment produced biomass that was significantly greater than (p *<* 0.05) the other treatments (significant differences among treatments are denoted by different letters). The bacterial abundance sample for the C:P = 100 treatment was lost before analysis and is therefore not reported in panel D.

### 3.2 PLANKTONIC AND BIOFILM COMMUNITY STRUCTURE

#### 3.2.1 Alpha diversity

We used rarefaction curves to evaluate alpha diversity in all treatments for both the plankton and the biofilm communities. Rarefaction curves showed heterotroph and photoautotroph OTU richness was consistently higher in the biofilm compared to the planktonic communities (Figure 3). For both the photoautotroph and heterotroph sequence datasets the biofilm and plankton communities had the fewest OTUs in the highest C treatment (C:P = 500) (Figure 3). When planktonic rRNA gene sequences from all planktonic samples were pooled, individual biofilm heterotroph community richness still generally exceeded the pooled planktonic heterotroph richness. For photoautotrophs, the individual biofilm richness was similar to the pooled photoautotroph planktonic richness(Figure 4).

**Figure 3.**
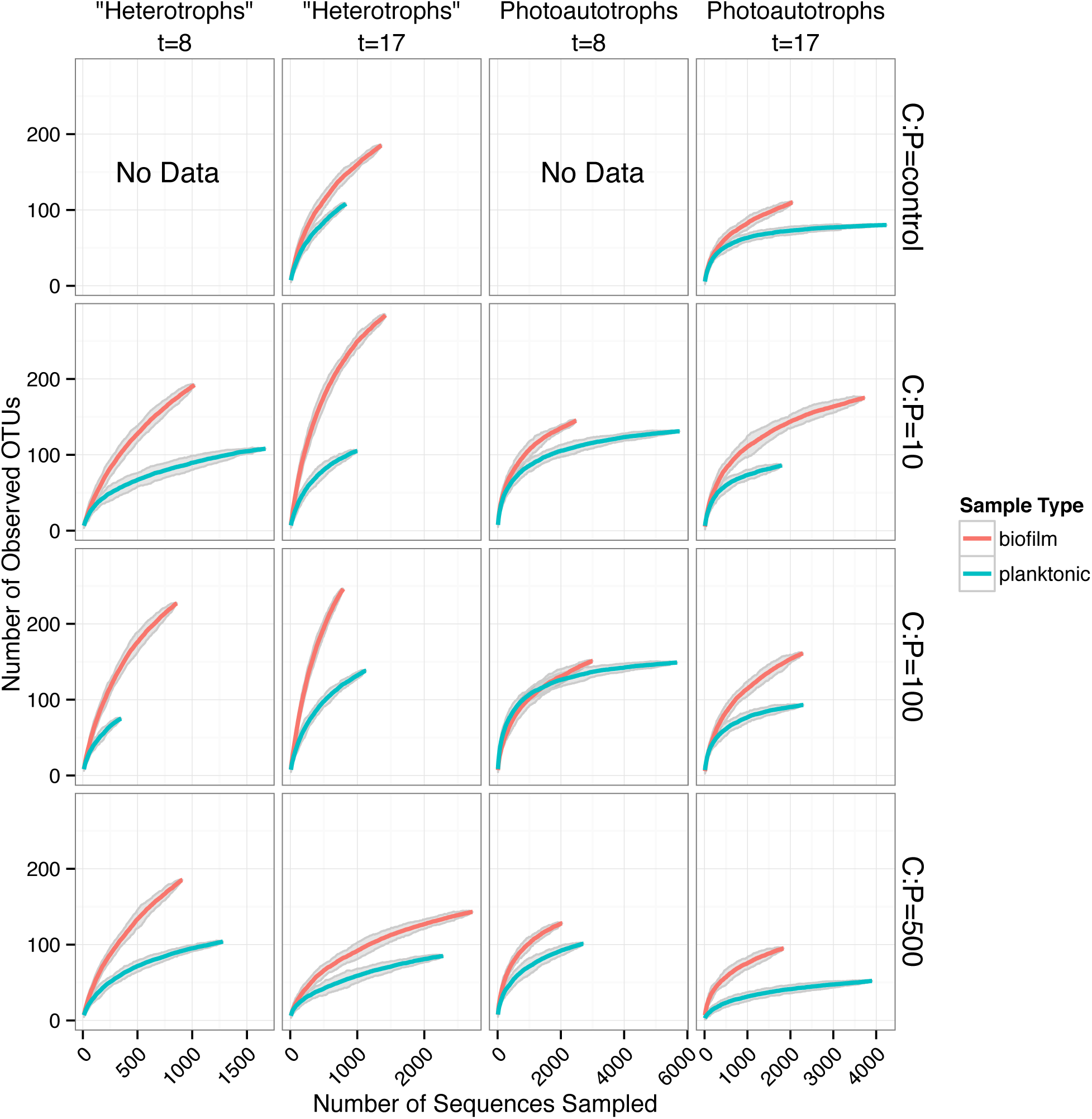
Rarefaction curves for all biofilm versus plankton libraries. Each panel represents a single C:P treatment and time point. Richness is greater for all biofilm communities when compared to corresponding planktonic communities. Gray ribbons are 99% confidence intervals around each rarefaction point based on variance from 25 re-samplings.

**Figure 4.**
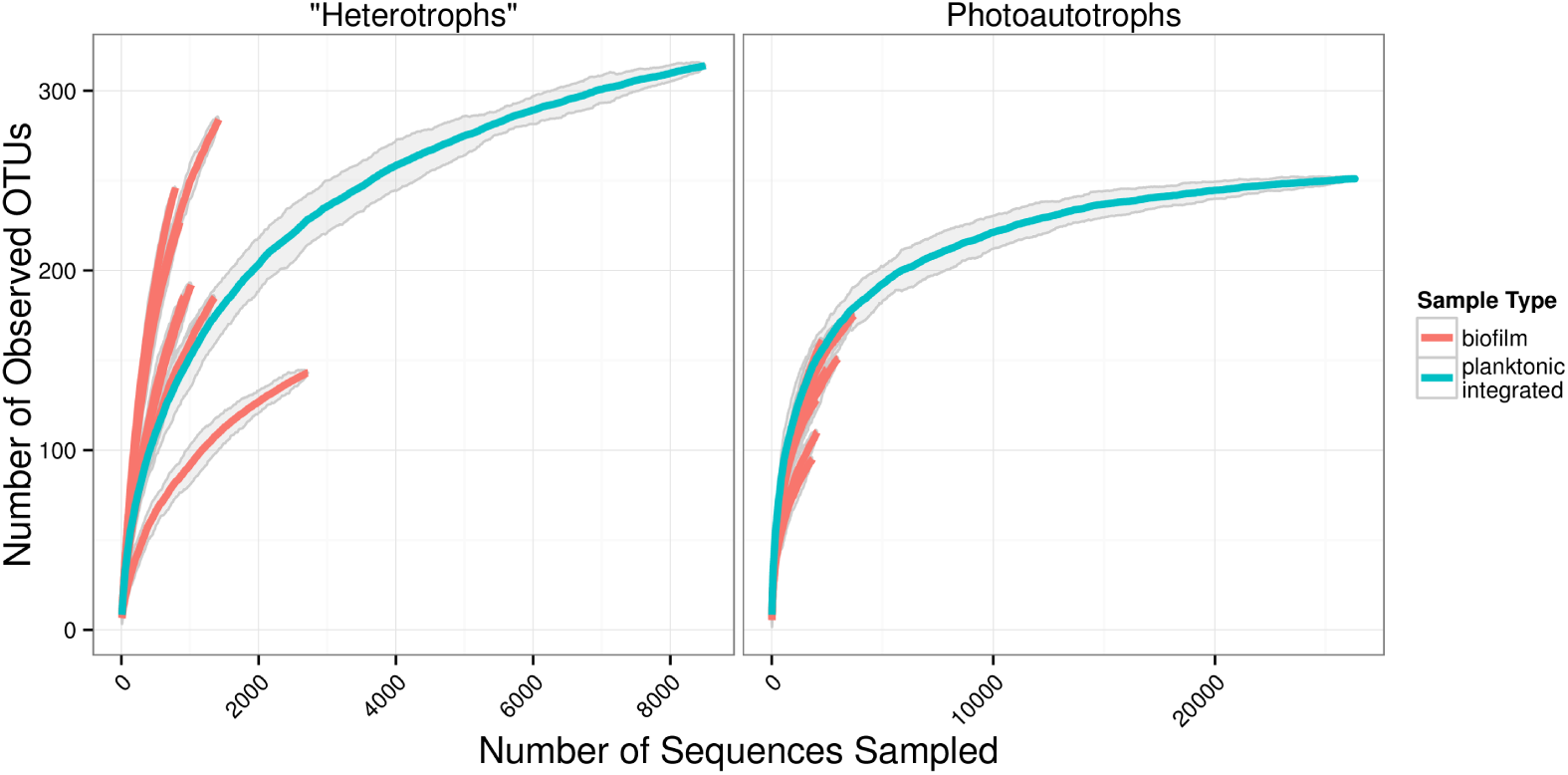
Rarefaction plots for all samples. Planktonic libraries have been integrated such that the count for each OTU is the sum of counts across all samples. Gray ribbons are 99% confidence intervals around each rarefaction point based on variance from 25 re-samplings.

#### 3.2.2 Community membership biofilm versus plankton

Heterotroph community membership between the plankton and biofilm communities was notably different for all treatments except for the highest C treatment where the plankton and biofilm communities during the second sampling event (day 17) were more similar to each other than any other community (Figure 5). Photoautotroph plankton and biofilm communities were also different in OTU composition, however, the similarity among photoautotroph plankton and biofilm communities in the highest C treatment was not observed as it was for the heterotroph communities (Figure 5).

**Figure 5.**
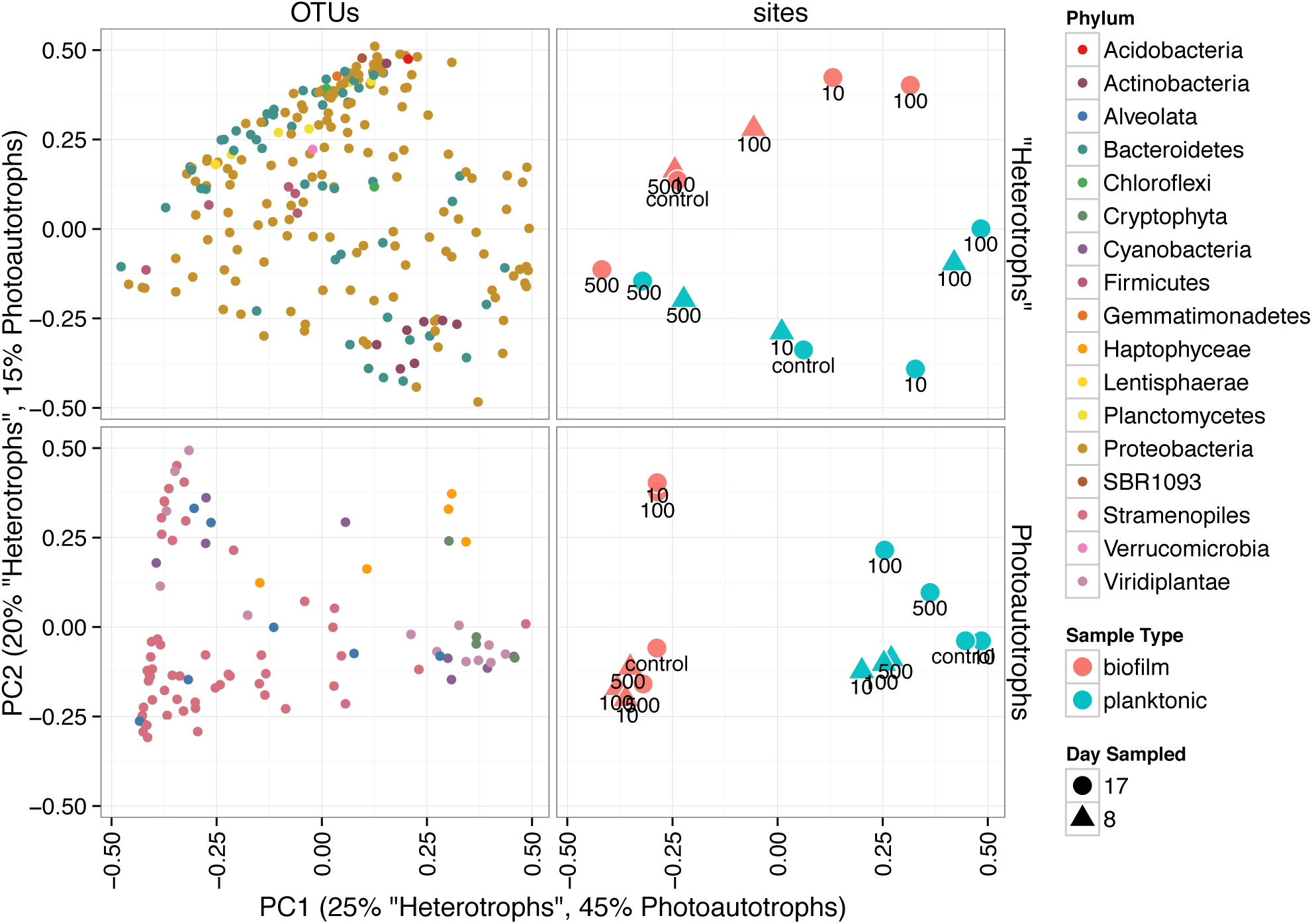
Principal coordinates ordination of bray-curtis distances for 23S rRNA plastid libraries and 16S rRNA gene libraries. OTU points are weighted principal coordinate averages (weights are relative abundance values in each sample) and the variance along each principal axis is expanded to match the site variance. Point annotations denote the amended C:P ratio for the mesocosm from which each sample was derived.

In heterotroph libraries, 19,978 sequences were distributed into 636 OTUs 58% of quality controlled sequences fell into the top 25 OTUs in order of decreasing sum of relative abundance across all samples. In photoautotroph libraries 44,719 23S plastid rRNA gene sequences were distributed into 359 OTUs 71% of sequences fell into the top 25 OTUs sorted by mean relative abundance across all samples.

To investigate differences in community structure and membership between the heterotroph biofilm and overlying planktonic communities we identified the most enriched OTUs in biofilm compared to the planktonic communities and vice versa. The most enriched OTUs were enriched in planktonic samples (with respect to biofilm) as opposed to biofilm samples (Figure 6). This is consistent with the higher alpha diversity in biofilm communities compared to planktonic communities and evidence that sequence counts were spread across a greater diversity of taxa in the biofilm libraries compared to the planktonic libraries (i.e. biofilm communities had higher evenness than planktonic communities). Of the top 5 enriched heterotroph OTUs between the two lifestyles (biofilm or plankton), 1 is annotated as *Bacteroidetes*, 1 *Gammaproteobacteria*, 1 *Betaproteobacteria*, 1 *Alphaproteobacteria* and 1 *Actinobacteria* and all 5 were enriched in the planktonic libraries relative to biofilm (Table 1). Of the 25 most enriched OTUs among lifestyles only 2 heterotroph OTU centroid DNA sequences shared high sequence identity (*>*= 97%) with cultured isolates (”OTU.32” and “OTU.48”, Table 1).

**Figure 6.**
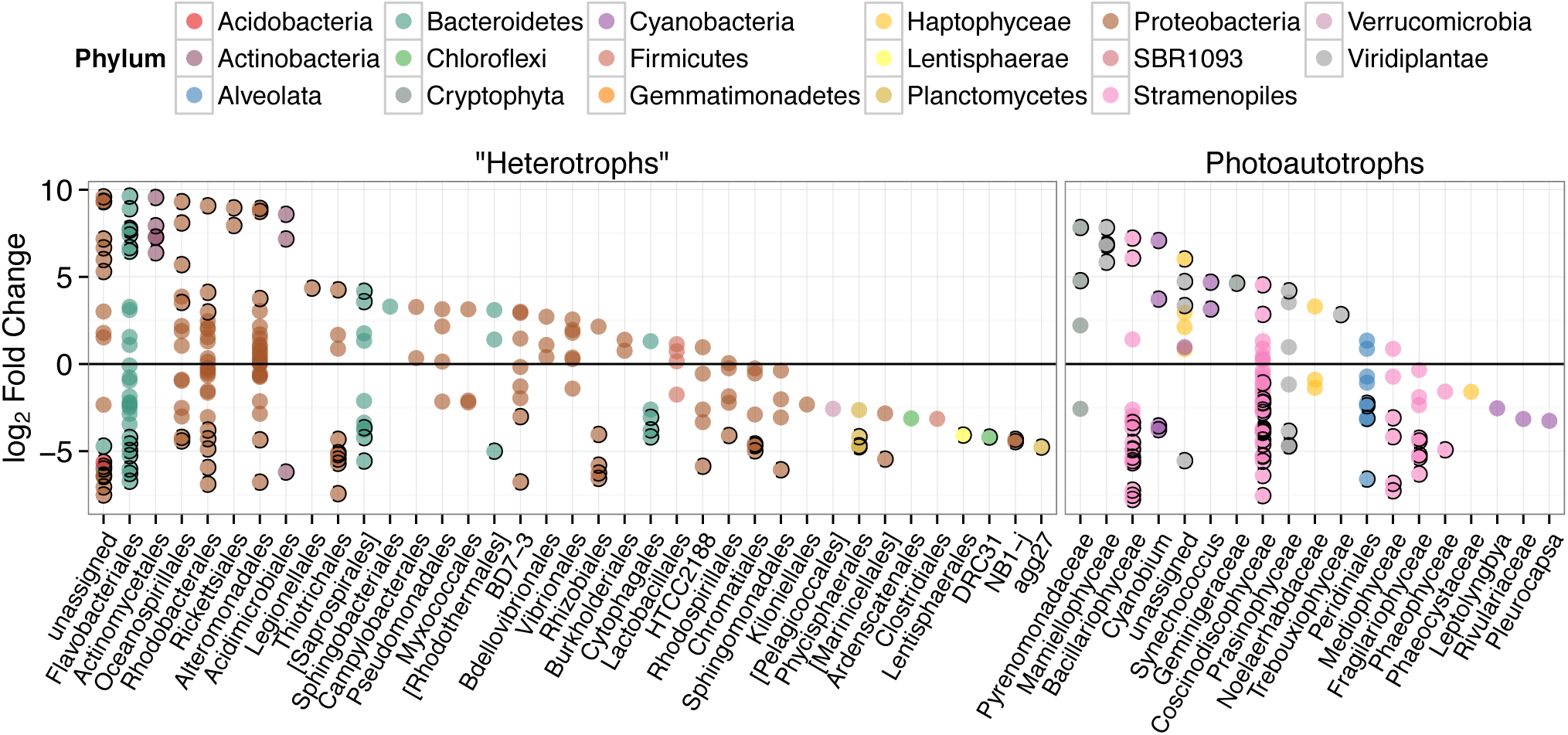
*log*_2_ of lifestyle OTU abundance fold change between biofilm and plankton communities. Each point represents one OTU and points are grouped along the x-axis by Order. Outlined points have adjusted p-values below a false discovery rate of 0.10. Positive fold change values represents enrichment in planktonic samples.

**Table 1.**
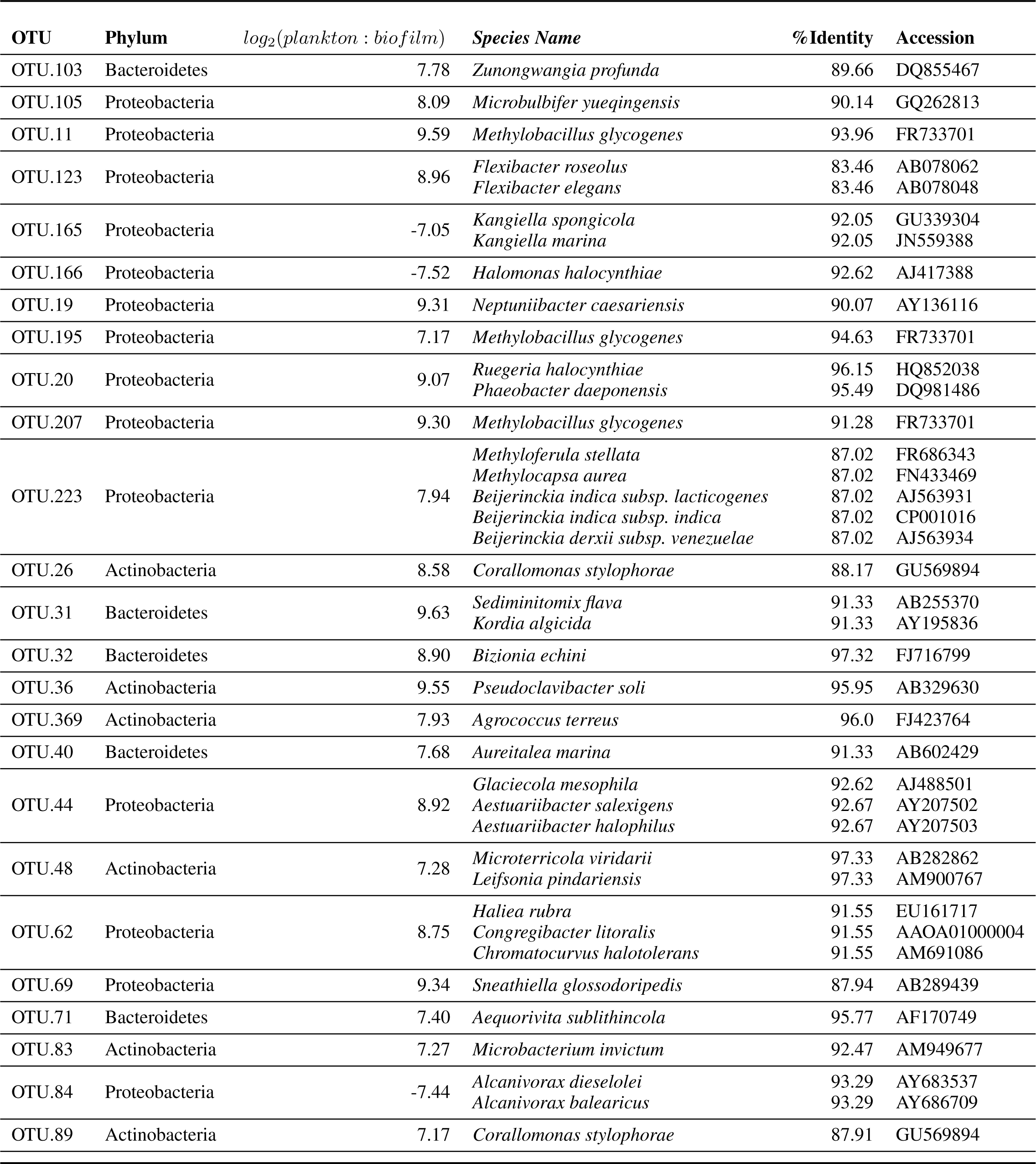
Results for BLAST search against Living Tree Project (top 25 lifestyle enriched bacterial OTUs (Operational Taxonomic Unit)

We similarly assessed membership among biofilm and plankton in the photoautotroph communities. Photoautotroph 23S plastid rRNA gene sequence libraries also clustered strongly by lifestyle (Figure 5). Biofilm libraries were predominantly enriched in *Stramenopile* OTUs whereas planktonic libraries were enriched in *Haptophyceae*, *Cryptophyta* and *Viridiplantae* OTUs based on OTU positions in sample ordination space (Figure 5, see Ordination Methods). When photoautotroph OTUs were ordered by differential abundance between lifestyles (see Figure 6), 16 of the 25 most enriched OTUs were enriched in the biofilm and 9 were enriched in the planktonic samples. Fourteen of these 16 biofilm enriched OTUs were *Stramenopiles* of class *Bacillarophyta*, the remaining OTUs were classified as members of the *Chlorophyta* and *Dinophyceae*. The 9 planktonic enriched OTUs (above) were distributed into the *Viridiplantae* (5 OTUs), *Cryptophyta* (1 OTUs), *Haptophyceae* (1 OTU), *Stramenopiles* (1 OTU) and cyanobacteria (1 OTU). The 10 most enriched photoautotroph OTUs between lifestyles were evenly split between planktonic and biofilm enriched OTUs. As with the heterotrophs, photoautotroph OTU fold change between lifestyles are qualitatively consistent with OTU positions in sample ordination space (see Figures 6 and 5).

In both the heterotroph and photoautotroph communities low abundance members of the planktonic communities became highly abundant members of the biofilm (Figure 7). The separation in community membership among biofilm and planktonic communities is supported statistically by the Adonis test (**Anderson**, 2001) for both the heterotroph and photoautotroph libraries (p-value 0.003 and 0.002, respectively). The lifestyle category represents 18% and 45% for pairwise sample distance variance in heterotroph and photoautotroph libraries, respectively. The Adonis test result is also consistent with lifestyle (biofilm versus planktonic) clustering along the first principal component for the photoautotroph libraries but not for the heterotroph libraries (Figure 5).

**Figure 7.**
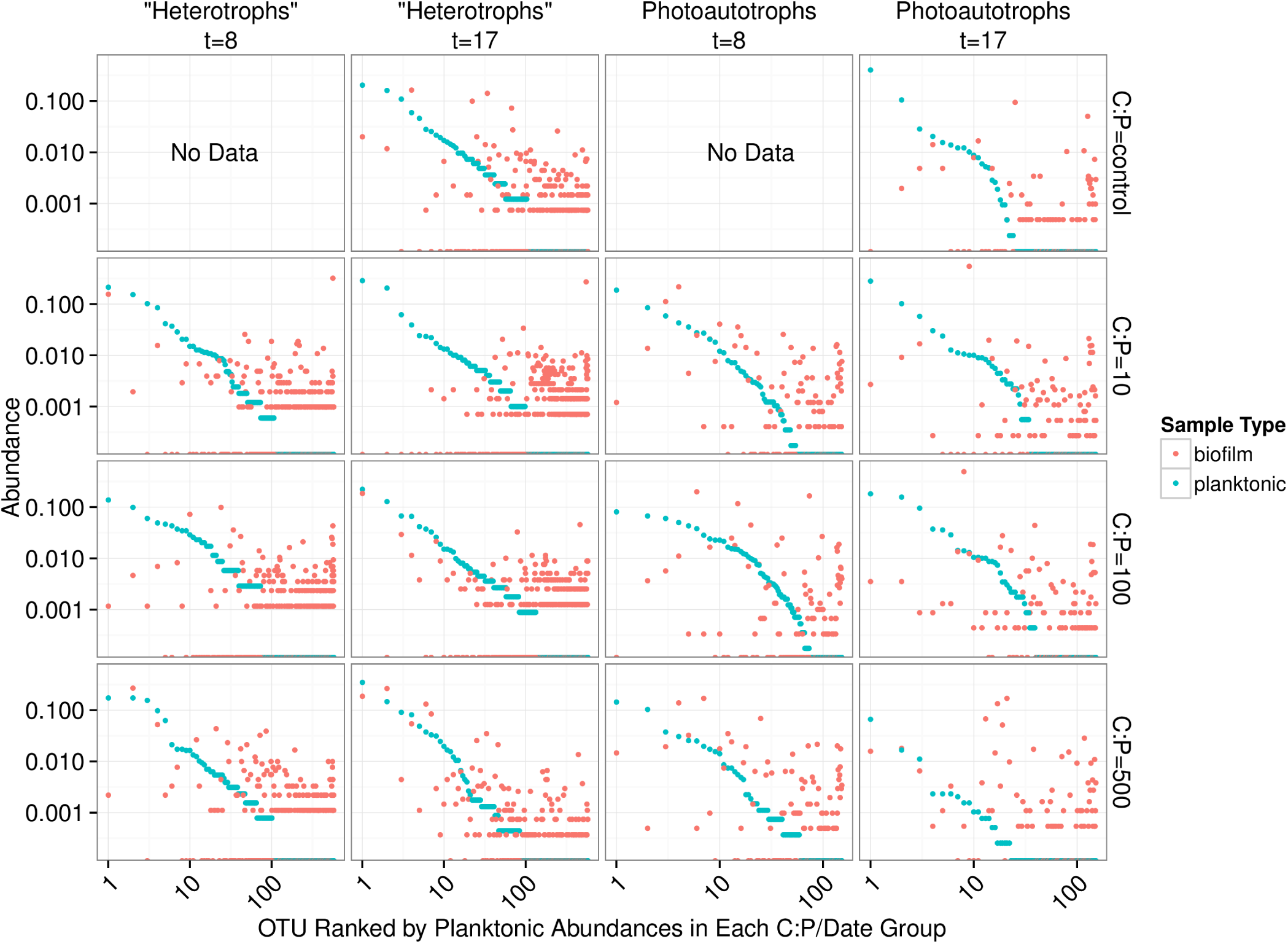
Rank abundance plots. Each panel represents a single time point and C:P. The “rank” of each OTU is based on planktonic sample relative abundance. Each position along the x-axis represents a single OTU. Both the x and y axes are scaled logarithmically.

#### 3.2.3 Heterotroph community membership changes with C amendments

Although community membership was predominately driven by lifestyle, we also investigated how resource amendments affected community membership and structure. Because the abiotic (e.g. DOC) and all biomass indicators (e.g. heterotroph abundance, Chl *a*) were only significantly different in the highest resource C:P treatment we compared resource C:P = 500 (high C) to all other mesocosms (i.e. control, C:P = 10 and C:P = 100 − low C). The high and low carbon amended mesocosms had statistically different heterotroph communities (Adonis p-value 0.018) but not photoautotroph communities (Adonis p-value 0.59). Nine heterotroph OTUs were enriched in the high C treatment relative to low C. Four of the 9 high C enriched OTUs were annotated as *Alteromonadales*, 3 as *Campylobacterales* and 1 each into *Vibrionales* and *Pseudomonadales*. The most enriched OTU in low C mesocosms was annotated as belonging to the “HTCC2188” candidate order and shared 99% identity with a 16S sequence annotated as “marine gamma proteobacterium HTCC2089” (accession AY386332). We only observed differences at the community level between high and low C amendments in the heterotroph communities and therefore did not assess differential abundance of OTUs between high and low C treatments in photoautotroph communities.

## 4 DISCUSSION

### 4.1 BIOMASS POOL SIZE

The goal of this study was to evaluate how changes in available C affected the biomass pool size and composition of planktonic and biofilm communities. Our results suggest that C subsidies increased heterotroph biomass in both plankton and biofilm communities. C amendments also resulted in decreased photoautotroph biomass in the plankton community, but there was no significant change in biofilm photoautotroph biomass between resource treatments. Although the DOC concentration in the highest C treatment was significantly higher than the other treatments, the concentrations we measured were in the range of those reported in natural marine ecosystems (**Mopper et al.**, 1980) and it is has been noted that glucose concentrations in coastal marine ecosystems may fluctuate over several orders of magnitude (**Alonso and Pernthaler**, 2006). The changes in the biomass pool size that did occur were consistent with changing relationships (commensal to competitive) between the autotrophic and heterotrophic components of the plankton communities but not necessarily of the biofilm communities. While we recognize that other mechanisms may drive the shift in biomass pool size of these two components of the microbial community (e.g. increased grazing pressure on the photoautotrophs with C additions, or production of secondary metabolites by the heterotrophs that inhibit algal growth) previous studies **Stets and Cotner** (2008a); **Cotner and Biddanda** (2002) and the data reported here suggest that altered nutrient competition among heterotrophic and photoautotrophic members of the plankton is the most parsimonious explanation for the observed shift in biomass pool size.

### 4.2 BIOFILM AND PLANKTON ALPHA AND BETA DIVERSITY

Beyond changes in the biomass pool size of each community, we explored how shifts in resource C affected the membership and structure of each community, and the recruitment of plankton during biofilm community assembly. Intuitively, shifts in planktonic community composition should alter the available pool of microorganisms that can be recruited into a biofilm. For example, if planktonic diversity increases, the number of potential taxa that can be recruited to the biofilm should also increase, potentially increasing diversity within the biofilm. Similarly, a decrease in mineral nutrients available to photoautotrophs should decrease photoautotroph pool size, potentially decreasing photoautotroph diversity and therefore candidate photoautotroph taxa that are available for biofilm formation. In addition, C in excess of resource requirements may increase the production of extracellular polysaccharides (EPS) by planktonic cells thus increasing the probability that planktonic cells are incorporated into a biofilm by adhesion. Each of these mechanisms suggest that an increase in labile C to the system should result in increased alpha diversity in heterotrophic plankton and heterotrophic biofilm communities while decreasing alpha diversity within both planktonic and biofilm photoautotroph communities.

We highlight three key results that we find important for understanding aquatic biofilm assembly. First, biofilm community richness exceeded planktonic community richness (Figure 3) in all mesocosms. Second, for the control, C:P = 10 and C:P = 100 resource treatments the membership and structure of the heterotroph biofilm and plankton communities were more similar within a lifestyle (plankton versus biofilm) than within a resource treatment. However, for the bacteria in the highest C treatment (C:P = 500) both membership and structure of biofilm and planktonic communities at day 17 were more similar to each other than to communities from other treatments (Figure 5). Third, C subsides acted differently on the photoautotroph and heterotroph communities. Specifically, while the highest level of C subsidies (C:P = 500) resulted in a merging of membership in the heterotroph plankton and heterotroph biofilm communities the same merging of membership was not observed for the photoautotroph biofilm and plankton communities which had distinct membership in all treatments.

We propose two potential mechanisms for the increased richness of the biofilm communities relative to the planktonic community richness. First, it is possible that the planktonic community composition of our flow through incubators was dynamic in time. In this case the biofilm community would represent a temporally integrated sample of the planktonic organisms moving through the reactor resulting in higher apparent alpha diversity (i.e. mass effects would be the dominant assembly mechanism). Second, the biofilm environment may disproportionately enrich the low abundance members of the planktonic community. In this case it is probable that the biofilm would incorporate the most abundant members from the planktonic community (i.e. mass effects) but also select and enrich (i.e. species sorting) the least abundant members of the planktonic community resulting in a higher level of detectable alpha diversity. The second mechanism would result if the biofilm environment represented a more diverse microhabitat including sharply delineated oxygen, nutrient and pH gradients that are not present in the planktonic environment. In this case the more diverse microhabitat would be able to support a more diverse community due to an abundance of additional environmental habitats (i.e. niches).

We evaluated the first mechanism by comparing membership among the plankton samples taken 9 days apart (day 8 and day 17). While heterotrophic plankton communities were not identical between the time points (Figure 5), communities within a treatment were more similar to each other than other heterotroph plankton communities regardless of time. In addition, the control and two lowest C treatments (C:P = 10 and C:P = 100) separated completely from biofilm communities in principle coordinate space (Bray-Curtis distance metric). This suggests that the biofilm community was not integrating variable bacterioplankton community membership, but rather was at least in part selecting for a community that was composed of distinct populations when compared to the most abundant members of the plankton community. As noted above, in the highest C treatment (C:P = 500) the heterotroph biofilm and plankton community membership had significant overlap at the final timepoint (Figure 5). However, heterotrophic plankton community composition for the highest C treatment among timepoints (8 and 17 days) were also qualitatively as similar to each other as any other community. Thus, variable planktonic community composition among timepoints would not explain the higher diversity observed in the biofilm compared to the planktonic community. Rather, two results point to enrichment of planktonic community members within the biofilm as the mechanism for higher diversity in the biofilm compared to the plankton. First, the increasing similarity between the plankton and the biofilm communities between each time point in the highest resource C treatment suggests that *in situ* resource conditions were sufficient to alter the relative abundance of the populations within each community. Second, an analysis of the OTU relative abundance in biofilm and planktonic libraries where OTUs are sorted by planktonic sample rank (Figure 7) shows that the least abundant members of the plankton community were routinely highly abundant within the biofilm community. This was true for both photoautotroph and heterotroph communities, at all treatment levels and both timepoints. While we did not (could not) specifically measure niche diversity within the biofilm communities our results suggest that the biofilm habitat selected for unique members of the photoautotroph and heterotrophic community that were in low abundance in the planktonic habitat but readily became major constituents of the biofilm community.

Few studies have simultaneously evaluated the relationship among membership and/or diversity of the plankton and the biofilm community from complex environmental microbial communities. One notable study looked at planktonic community composition and biofilm formation on glass beads placed for three weeks in three boreal freshwater streams (**Besemer et al.**, 2012). While that study system is markedly different than our study, the analyses and questions addressed in each study were sufficiently similar to merit comparison. **Besemer et al.** (2012) concluded that the biofilm community membership was most likely driven by species sorting over mass effects. This is consistent with what we report here. However, in the **Besemer et al.** (2012) study the authors reported that planktonic diversity was significantly higher relative to biofilm diversity (the opposite of what we found in our study). Given the differences in the source of the planktonic community among studies, this result is not surprising. While biofilm communities were established on glass beads in **Besemer et al.** (2012) and glass slides (this study) over a similar time period (21 days, **Besemer et al.** (2012) and 17 days this study) the origin of the planktonic community in each study was different. The **Besemer et al.** (2012) study was conducted in three boreal streams during snow melt when connectivity between the terrestrial and aquatic habitats was high and potentially highly variable depending on how hydrologic pathways differed among precipitation events. In this study the source community was a marine intake located approximately 200 m from the shore during July when communities are more stable over the 17 day period of the incubation. A separate study conducted in alpine and sub-alpine streams clearly showed that stream plankton communities reflected localized precipitation events and could be traced largely to soil source communities from drainages within the watershed (**Portillo et al.**, 2012). While planktonic communities in lake ecosystems can be linked to soil communities in the watershed, as residence time of the system slows the relative influence of species sorting increases. Thus, in headwater ecosystems stream plankton communities can often be composed primarily of soil organisms (**Crump et al.**, 2012). In addition to the diverse source communities the **Besemer et al.** (2012) study sampled the plankton community at multiple timepoints and integrated the samples before sequencing further increasing community richness as compared to the current study where the plankton community was sampled and analyzed only at two independent timepoints. Indeed, when we pool OTU counts from all planktonic libraries and compare the rarefaction curve of the pooled planktonic libraries (photoautotrophs and heterotrophs) against sample-wise biofilm libraries, we found more total heterotroph and photoautotroph planktonic OTUs than in any given single biofilm sample. It appears, however, that sample-wise heterotroph biofilm rarefaction curves may exceed the integrated planktonic curve upon extrapolation and most exceed the integrated planktonic curve at sampling depths where data is present for the biofilm and pooled planktonic library (Figure 4). This result is consistent with our conclusion that temporal heterogeneity in the plankton was not sufficient to explain the higher diversity in the biofilm sample but would explain the relative differences between planktonic and biofilm diversity found in **Besemer et al.** (2012) compared to this study.

In addition, for this study, it is important to note that biofilm community richness peaked at the intermediate treatment (C:P = 100) and appeared to decrease between each time point although with only two time points it was unclear how pronounced the temporal effect was nor is it possible assess the statistical significance of this effect (Figure 3). Since biomass of the plankton and the biofilm increased with increasing C subsidies the intermediate peak in OTU richness is consistent with a classic productivity-diversity relationship that has been shown for many ecosystems and communities both microbial and otherwise. However, as with other experiments our experimental design did not allow us to tell whether resources drove productivity that subsequently drove changes in diversity or whether resources drove diversity which altered productivity. Rather, we note that as diversity decreased in the highest C treatment, heterotrophic plankton and biofilm membership became increasingly similar. This suggests that environments that contained high amounts of labile C selected for fewer dominant taxa, overwhelming the lifestyle species sorting mechanisms that appeared to dominate biofilm community assembly in all other treatments. Similarly, while we did not measure extracellular polymeric substances (EPS), direct microscopy showed that planktonic cells in the highest C treatment (C:P = 500) were surrounded by what appeared to be EPS. Because biofilm EPS also appeared to increase from the low to high C treatments it is possible that abundant planktonic cells were more readily incorporated into biofilms due both to increased “stickiness” of the planktonic cells as well as the biofilm itself. While we did not observe flocculating DOC which has been shown to dominate high DOC environments in nature, we did measure a substantial increase in DOC in the C:P = 500 treatment which was more than 2-fold higher than any of the other treatments. Thus additional adhesion of the plankton and the biofilm may also explain the merging of the planktonic and biofilm heterotroph membership in the highest C treatment.

### 4.3 LIFESTYLE (BIOFILM OR PLANKTONIC) ENRICHED OTUS

There are only a few studies that attempt to compare biofilm community composition and the overlying planktonic community (**Besemer et al.**, 2007, 2012; **Jackson et al.**, 2001; **Lyautey et al.**, 2005). Those studies illustrate community composition among the two habitats are unique, with few taxa found in both. This is consistent with our findings in this experimental system with a natural marine planktonic source community. In addition, our study also evaluated photoautotroph community composition which showed a similar result suggesting that both the photoautotroph and heterotroph biofilm communities are comprised of phylogenetically distinct organisms that exist in low abundance in the surrounding habitat (i.e. the plankton) but are readily enriched in the biofilm lifestyle. Most of the biofilm enriched photoautotroph OTUs were *Bacillariophyta* although there were also many *Bacillariophya* OTUs enriched in the planktonic libraries. We also found more *Cryptophyta* and *Viridiplantae* were enriched in the planktonic photoautotroph libraries. It appears that these broad taxonomic groups were selected against in biofilms under our experimental conditions. Heterotroph OTUs enriched in planktonic samples displayed more dramatic differential abundance patterns than heterotroph OTUs enriched in biofilm samples, but, biofilm enriched heterotroph OTUs were spread across a greater phylogenetic breadth (Figure 6). This is also consistent with the idea of greater niche diversity in the biofilm environment as opposed to the plankton. Greater niche diversity should select for a more diverse set of taxa but individual taxa would not be as numerically dominant in a more uniform environment like the planktonic environment. At the Order level, enriched heterotroph OTUs tended to have members that were enriched in both the plankton and the biofilm suggesting the phylogenetic coherence of lifestyle is not captured at the level of Order. It should be noted, however, that taxonomic annotations in reference databases and therefore environmental sequence collections show little equivalency in phylogenetic breadth between groups at the same taxonomic rank (**Schloss and Westcott**, 2011). Unfortunately, at higher taxonomic resolution (e.g. Genus-level), groups did not possess a sufficient numbers of OTUs to evaluate coherence between taxonomic annotation and lifestyle. Carbon amendments did not affect photoautotroph library membership and structure to the same degree as it affected heterotroph library composition. As expected, heterotroph OTUs enriched in the high C amended mesocosm (C:P = 500) include OTUs in classic copiotroph families such as *Altermonodales* and *Pseudomonadaceae*. Interestingly, the most depleted OTU in the high C treatments is annotated as being in the HTCC2188 order of the *Gammaproteobacteria* and shares 99% sequence identity with another “HTCC” strain (accession AY386332). HTCC stands for ‘high throughput culture collection’ and is a prefix for strains cultured under low nutrient conditions (**Cho and Giovannoni**, 2004; **Connon and Giovannoni**, 2002).

### 4.4 CONCLUSION

In summary this study shows that changes in low resolution community level dynamics are concurrent with changes in the underlying constituent populations that compose them. We found that autotrophic pools and heterotrophic pools responded differently to amendments of labile C as hypothesized. Notably while C amendments altered both pool size and membership of the heterotroph communities we did not see similar dynamics within the photoautotroph communities. Planktonic photoautotrophs decreased in response to C amendments presumably in response to increased competition for mineral nutrients from a larger heterotroph community, however there was not a similar decrease in biofilm photoautotroph community. In addition membership of the photoautotroph communities between the plankton and biofilm lifestyles did not become more similar in the photoautotrophs as it did for the bacterial heterotrophs in the highest C treatment. Consistent with a growing body of work our results suggest that complex environmental biofilms are a unique microbial community that form from taxa that are found in low abundance in the neighboring communities. This membership was affected by C amendments for heterotrophic but not photoautotrophic microbes and then only in the most extreme resource environment. This suggests that lifestyle is a major division among environmental microorganisms and although biofilm forming microbes must travel in planktonic form at some point, reproductive success and metabolic contributions to biogeochemical processes comes from those taxa primarily if not exclusively while they are part of a biofilm. Our results point to lifestyle (planktonic or biofilm) as an important trait that explains a portion of the exceptional diversity found in snapshots used to characterize environmental microbial communities in space and time.

## 5 ACKNOWLEDGEMENTS

This research was conducted as part of the 2010 Marine Biological Laboratory’s Microbial Diversity Course. Funding was supplied by the MBL, the Bernard Davis Endowed Scholarship Fund, and the Selman A. Waksman Endowed Scholarship. We would like to thank Dan Buckley and Steve Zinder for organizing the course, Marshall Otter, Mathew Erickson, Hugh Ducklow and Jay T. Lennon for analytical support and the Austrian FWF MICDIF award to Tom Battin for salary support.

